# Identification of drug candidates against glioblastoma with machine learning and high-throughput screening of heterogeneous cellular models

**DOI:** 10.1101/2025.03.06.641926

**Authors:** Vanessa Smer-Barreto, Richard J. R. Elliott, John C. Dawson, Álvaro Lorente-Macías, Asier Unciti-Broceta, Diego A. Oyarzún, Neil O. Carragher

## Abstract

Glioblastoma multiforme (GBM) is an aggressive primary brain tumour that presents significant treatment challenges due to its complex pathology and heterogeneity. The lack of validated molecular targets is a major obstacle for discovering new therapeutic candidates, with no new effective GBM therapies delivered to patients in over two decades. Here, we report the identification of compounds that target the GBM stem cell survival phenotype. Our approach employs machine learning (ML) predictors of cell survival trained on high-throughput, image-based, phenotypic screening data for 3,561 compounds, at multiple concentrations, across a panel of six heterogeneous, patient-derived, GBM stem cell lines. We computationally screened more than 12,000 compounds spanning various chemical classes. Experimental validation of ML-identified candidates across the GBM stem cell lines led to the identification of three compounds with activity against the GBM phenotype. Notably, one of our validated hits, the Hsp90 inhibitor XL888, displayed targeted elimination of all six GBM stem cell lines with IC50 in the nanomolar range. The other two compounds, which displayed broad activity across multiple GBM cell lines with distinct cell line sensitivities, offer routes for future personalised medicine campaigns. Our work demonstrates the use of phenotypic screening in tandem with ML can effectively identify therapeutic leads for personalised treatments in highly heterogeneous indications with few known molecular targets.

## Introduction

Glioblastoma multiforme (GBM) is the most common and aggressive primary brain tumour found in human adults and is characterised by substantial heterogeneity of genetic drivers and the tumour microenvironment^1–3^. Patients have poor prognosis and limited treatment options (typically surgery followed by chemoradiation) that lead to the emergence of resistance. For the past 20 years, the standard of care for newly diagnosed GBM patients has consisted of surgery, temozolomide (TMZ), and ionizing radiation (IR) prolonging the median overall survival of patients from 12 to 15 months^4,5^. Large-scale genomic analyses have enhanced our understanding of the molecular biology of GBM, which has supported the classification of GBM into various subtypes^6–8^. However, this new understanding of GBM has not yet delivered new effective treatment strategies. Current trials are limited to narrow subtypes of disease characterised by well established druggable target pathways that have failed to demonstrate durable responses^9,10^.

The dismal prognosis of this cancer of unmet need urgently calls for innovative approaches to find early drug candidates that can overcome the challenges of heterogeneity, including GBM stem cell plasticity and drug resistance, as well as effectively permeate the blood-brain-barrier. The intratumoural heterogeneity and ability of GBM stem cells to rapidly rewire signalling networks in response to inhibition of a single pathway^11^ confounds modern target-based drug discovery strategies. As a result, several studies have explored the use of target-agnostic phenotypic screening in GBM models, with particular efforts on the application of patient-derived cell lines^12–14^, drug repurposing^1,15^, and droplet microarray platforms^16^.

Computational screens based on Artificial Intelligence (AI) have recently gained notoriety due to their ability to sieve through vast collections of chemical data at scale, detecting patterns to identify compounds with a high probability of displaying a therapeutic phenotype of interest^17,18^. Machine learning models trained on phenotypic data have delivered a number of recent successes, including the discovery of senolytics^19^, novel antibiotics^20^, immune modulators^21^, and anti-inflammatory leads^22^, and constitutes one of the most exciting avenues for early drug discovery applications. The effectiveness of phenotypic- and AI-driven drug discovery depends on the availability of high-quality data acquired on disease-relevant model systems. In the case of GBM, it is possible to culture and propagate patient-derived GBM stem cells under conditions that maintain their stem cell like properties, thus recapitulating the heterogeneity and drug resistance mechanisms that lead to relapse in patients^23^.

In this work, we employed high-throughput phenotypic screening data and ML for discovering small molecule leads that hold potential for the development of alternatives to TMZ treatment of GBM. Using an in-house high-throughput screening assay of six patient-derived GBM stem cell lines^24^, we trained ML algorithms and successfully validated three predicted hits in cell lines representing the three main transcriptomic subtypes of GBM (proneural, classical, and mesenchymal). Medicinal chemistry evaluation identified one of these hits, XL888, as a promising chemical starting point for further novel drug design targeting inhibition of GBM stem cell growth and survival signalling.

Artificial intelligence and ML have been successfully applied in the context of GBM for drug repurposing^1^, tumour identification^25–27^, patient survival prediction^28–30^, and biomarker prediction^31,32^. To the best of our knowledge, our study is the first reporting the AI-powered discovery of chemical leads against the highly heterogeneous GBM phenotype. This demonstrates the power of data-driven methods in indications with poorly understood target biology, and highlights the benefits of employing training data that captures the heterogeneity of the disease of interest.

## Results

### High-throughput glioblastoma screen

The screening protocol across a panel of patient derived GBM stem cells is represented in Fig. 1a, with further detail provided in the Methods section and in Elliott et al^24^. We employed the ImageXpress-confocal Ht.ai high content imaging platform to quantify cell nuclei counts following small-molecule compound treatments as a readout for cell viability across six separate fields-of-view for each compound treatment. We performed our assay using six genetically diverse patient-derived cell lines GCGR-E13, GCGR-E21, GCGR-E28, GCGR-E31, GCGR-E34, and GCGR-E57, two from each of the most prominent subtypes of glioblastoma: classical, proneural, and mesenchymal (Fig. 1b). Our screen included a total of 3,561 compounds at variable concentrations (Fig. 1a) drawn from the Comprehensive anti-Cancer small Compound Library^33^ (C3L), the Kinase Chemogenomic Set (KCGS), the Library of Pharmacologically Active Compounds (LOPAC), the Prestwick chemical library, and the Targetmol anticancer library. We employed DMSO (0.1% v/v) as negative control and staurosporine (STS) and/or paclitaxel (PAC) as positive controls at concentrations of 1 μM STS (C3L, LOPAC, Prestwick chemical library), 0.1 μM PAC (KCGS) and both 5 nM PAC and 0.1 μM STS (Targetmol anticancer library).

**Figure 1.**
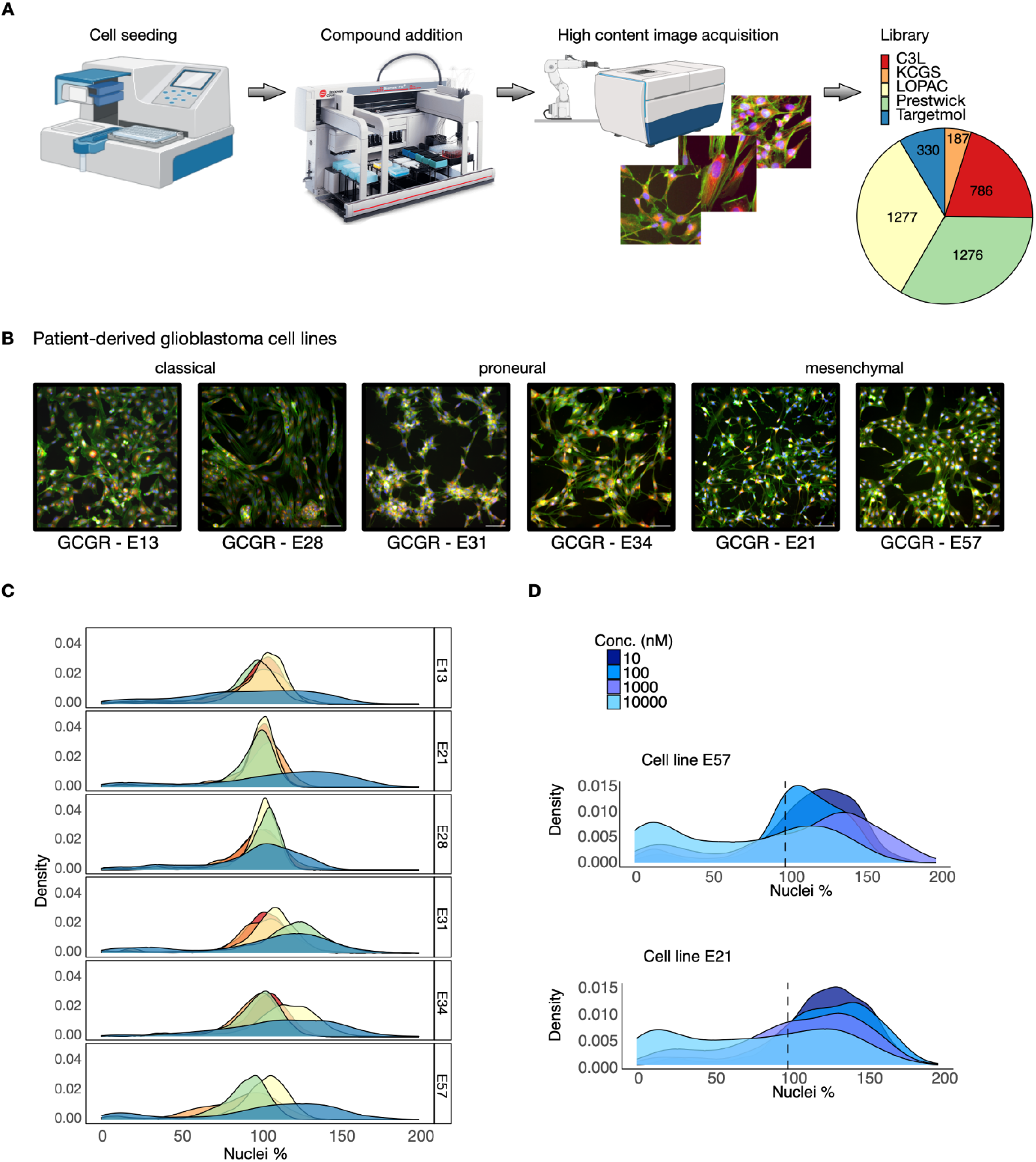
High-throughput phenotypic assay screening of heterogeneous, patient-derived GBM cell lines. (**A**) High content phenotypic assay screening workflow on GBM cells^24^. Five libraries were screened at multiple concentrations (C3L, 786 compounds at [0.01, 0.1, 1, 10] μm; KCGS, 187 compounds at [1, 10] μm; LOPAC, 1,277 compounds at [0.5, 3] μm; Prestwick, 1,276 compounds at [1, 10] μm; Targetmol-330, 330 compounds at [0.01, 0.1, 1, 10] μm). (**B**) Representative images of multiplex Cell Painting assay screen performed across of the three main GBM cellular subtypes: classical (GCGR-E13, GCGR-E28), proneural (GCGR-E31, GCGR-E34), and mesenchymal (GCGR-E21, GCGR-E57). The Cell Painting stains shown are Hoechst (blue), Phalloidin/WGA (green), and Endoplasmic reticulum (red). (**C**) Density plots over all six GBM cell lines of the percentage of nuclei counts after compound treatment with respect to the average DMSO negative control (100%) per plate. After removal of duplicates, the dataset contains 3,561 unique compounds. Colours correspond to each library (panel A), and only the highest concentration per library is shown. Cell line sensitivity to the compounds is variable across the cell lines; the bimodality displayed by the Targetmol library reveals a larger fraction of compounds that affect cell viability as compared to the other four libraries. **(D)** Density plots of the Targetmol library screen at all concentrations of cell line GCGR-E57 (top) and of cell line GCGR-E21 (bottom). The black dotted line represents the average DMSO negative control nuclei count. As the concentration increases, a larger number of compounds affects cell viability.

We observed variations in the proportion of compounds that affected target cell viability, both across cell lines and compound libraries, and this variation appeared to increase at higher concentrations (Fig. 1c). The density of compounds’ cell survival against DMSO control at the maximum concentration per library (C3L - 10uM, KCGS - 10uM, LOPAC - 3uM, Prestwick - 3uM, and Targetmol - 10uM) shows a bimodal distribution for the Targetmol library, with more treatments that affect the cells’ viability. In contrast, the LOPAC library’s unimodal distribution is centred on the DMSO control value of 100%. We further observed a higher susceptibility in some cell lines, as is the case for GCGR-E57, which displays a larger bimodal effect as the concentration increases on the Targetmol library screen (Fig. 1d top panel) in comparison to GCGR-E21 for the same conditions (Fig. 1d bottom panel).

### Machine learning models of GBM cell viability

We employed our screening data to train machine learning models of GBM cell viability in response to compound treatment, using a binary classification approach. Since our screen was performed at multiple concentrations and these varied across libraries, we labelled compounds as positive (leading to GBM cell death) or negative using a bespoke concentration-sensitive approach. This allowed us to make maximal use of the compound data and integrate all compounds into a single labelled dataset for model training. To this end, for each compound we first chose the concentration for which the phenotypic response was closest to 50% of nuclei count. Compounds were then labelled as positive if they eliminated at least two and not all six GBM cell lines using a cutoff of 35% nuclei count survival with respect to negative, DMSO controls. This strategy was designed to mitigate bias toward general cytotoxic compounds with a narrow therapeutic index. Details on our compound labelling pipeline can be found in the Methods.

For model training, we featurised each compound with physicochemical descriptors calculated with RDKit^34^, which have been successfully employed in previous models trained on phenotypic data ^19,35^. The resulting dataset contains 103 positive and 3,458 negative compounds for training, and is well distributed in the physicochemical feature space (Fig. 2a); low-dimensional representations using the UMAP algorithm^36^ do not suggest a per-library bias (Fig. 2a, left) and a suitable degree of chemical diversity for model training (Fig. 2a right). The distribution of pairwise Tanimoto distances (Fig. 2c) within positive and negative compounds suggests appropriate chemical diversity and reinforces the qualitative observations from the UMAP representations (Fig. 2a).

**Figure 2.**
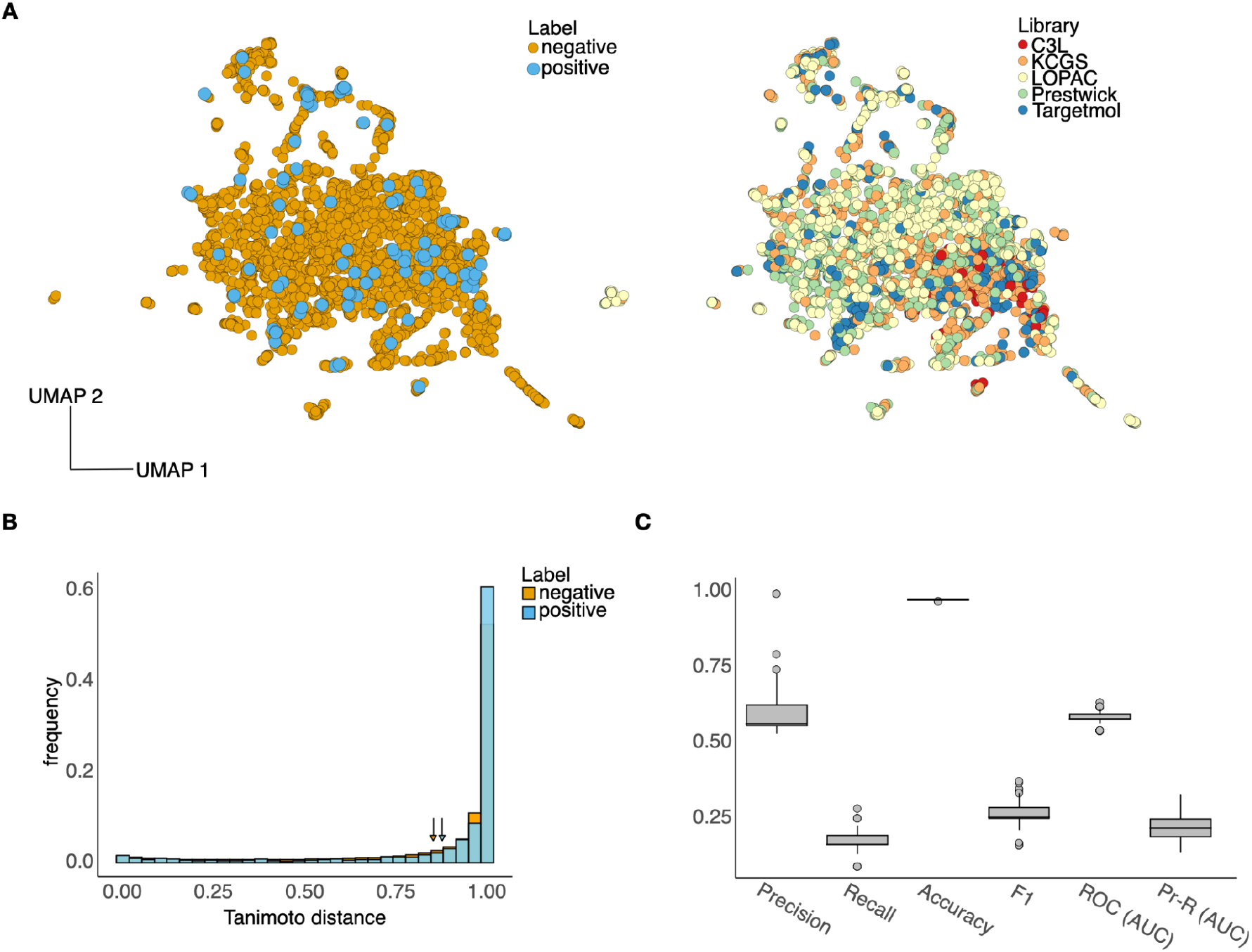
Visualisation of chemical space and training of machine learning models. (**A**) Two-dimensional UMAP visualisation of the 3,561 compounds employed for training machine learning models. Compounds were represented with a feature vector of 200 physicochemical descriptors computed with the RDKit^34^ software. UMAP plots were generated with number of neighbors 50, minimum distance 0.1, and spread 1. To train binary classifiers, the dataset was labelled as positives (compounds that eliminated 35% or more of the cells in the wells, and that did so for more than two but not all sixe cell lines) and negatives (all other compounds). Compounds were coloured according to their binary labels (left) and library of origin (right). (**B**) Tanimoto distances between compounds in the training set, split according to positives (N=10,506, all pairs) and negatives (N=11,954,298 pairs). The average Tanimoto distance across the positives is 0.88 ± 0.25; for the negatives cohort is 0.85 ± 0.26, suggesting suitable chemical diversity in both compound classes. Tanimoto distances were computed on RDKit features after normalization to zero mean and unit variance. **(C)** Performance metrics of 100 XGBoost algorithms trained on Monte Carlo re-sampled sets of 70% of compounds and tested on the remaining 30% (see Methods). Performance metrics include precision, recall, accuracy, F1 score, and area under the curve (AUC) of both Receiver Operating Characteristic and Precision-Recall curves. Box-and-whisker plots show the interquartile range and outliers across the 100 models, all of which surpass the naive accuracy baseline given by the class imbalance (2.9%).

Given the strong class imbalance in the data (2.9%), we devised a strategy to maximise the use of positive samples and, at the same time, yield confident predictions for downstream hit validation. To this end, we employed a Monte Carlo approach whereby a binary classifier was trained and tested multiple times with stratified resampled training (70%) and test data (30%). As a base model we chose an XGBoost classifier, which produces predictions using an ensemble of decision trees. After performing feature selection and Monte Carlo retraining 100 times (see Methods), we selected those models with test accuracy above the naive baseline, whereby all test compounds are called as the majority class (negative). Performance metrics of the selected models (Fig. 2c) show strong tradeoffs between precision (fraction of true positives out of all positive predictions) and recall (fraction of correct positive predictions), as commonly encountered in highly imbalanced classification tasks. Our strategy led to a precision of 0.6 ± 0.08 across N=100 models, which we deemed suitable for reducing the number of false positives for subsequent hit validation.

### Computational screen and hit validation against the GBM phenotype

We employed the ensemble of machine learning models for virtual screening and experimental testing of the predicted hits. For screening, we aggregated the anticancer Targetmol-3000 and Bioascent libraries. These libraries were chosen because the Targetmol library represents a collection of structurally diverse, medicinally active, and cell permeable FDA approved and clinical stage drug candidates each associated with rich documentation on structure, target, IC50 value and biological activity description. The Bioascent library contains broader structural diversity of an accessible 120,000 parent library of lead-like small molecules which are commercially available for follow up studies. After removal of duplicates and compounds present in the training data, we obtained a total 12,888 unique compounds for our screening library, with a good diversity with respect to the positives in the training data (Fig. 3a).

**Figure 3.**
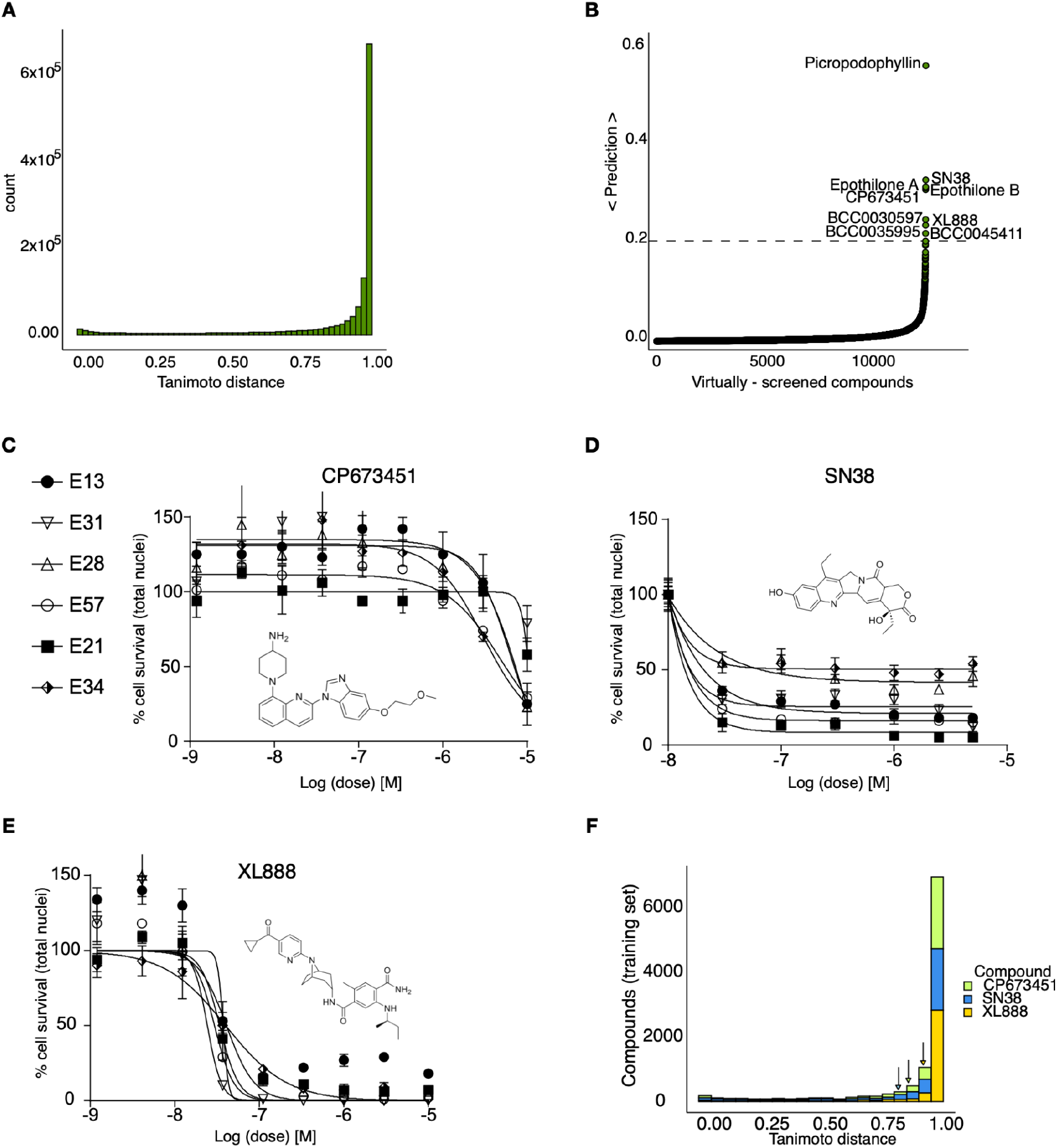
Computational screening and hit validation in GBM cellular models. (**A**) Compounds from two libraries, Targetmol-3000 and BioAscent, were selected for computational screening, giving a total of 12,888 unique compounds. The histogram shows the Tanimoto distance from each computationally-screened compound to each of the 103 positives in the training set; average Tanimoto distance is 0.86 ± 0.25 suggesting good diversity with respect to positive in training. (**B**) Computational screen of 12,888 compounds from the Targetmol-3000 and BioAscent libraries. Histogram shows the predicted probability of each compound affecting viability of GBM cellular models. The predictions were calculated for each of the 100 XGBoost models; histogram shows average prediction score for each compound across the trained models. Nine compounds scored above a 20% cut-off. Three compounds were not selected for hit validation because of closely-related analogues in the training set. Six compounds were taken forward for experimental validation in all six GBM cells lines. (**C-E**) Dose-response curves of three validated hits that affect the cell viability of GBM cell lines. Plots show dose-response curves and chemical structures of **(C)** CP673451, **(D)** SN38, and **(E)** XL888. SN38 and XL888 are effective at the nanomolar concentration range, while CP673451 is active at micromolar concentrations. Mean ± s.d. are shown from *n* = 3 experiments. (**F**) Stacked histograms of the Tanimoto distance between the three validated hits and each of the 3,561 compounds in the training set; average Tanimoto distance was 0.88 ± 0.24 for CP673451, 0.83 ± 0.28 for SN38, and 0.94 ± 0.17 for XL888, indicating substantial physicochemical differences between the hits and the compounds for training.

After querying the ensemble of machine learning models, we obtained highly selective prediction scores (Fig. 3b), with only nine predicted hits (0.07%) with an average score above 20%. This high selectivity is two orders of magnitude below the class imbalance of the training data and adds confidence in the model predictions. Examination of the predicted hits revealed that three compounds (epothilone A, epothilone B, and picropodophyllin) have close analogues in the training data (ixabepilone and podophyllotoxin), and thus were deliberately excluded from experimental testing. We tested the remaining six predicted hits in our GBM cell lines with a similar assay as for our high-content screening (Fig. 1a). Among the tested hits, three compounds (SN38, CP673451, and XL888) affected the viability of GBM cell lines (Fig. 3c-e, Supplementary Fig. 1): SN38 is the active metabolite of irinotecan, an approved topoisomerase inhibitor; CP673451 is a potent and selective inhibitor of platelet derived growth factor (PDGFR); XL888 is a known inhibitor of Heat shock protein 90 (Hsp90). The remaining three tested compounds (BCC0030597, BCC0035995, BCC0045411) did not impact GBM cell viability. The validated compounds displayed varied activity against the GBM phenotype (Table 1) with XL888 and SN38 having IC50 below 10uM for all six cell lines, while CP673451 only for a subset of them, possibly due to genetic heterogeneity conferring distinct sensitivity between cell lines (Fig. 3c-e).

**Table 1.** IC50 values (nM) of the three validated hits for each of the six GBM cell lines tested.

|  | classical |  | proneural |  | mesenchymal |  |
| --- | --- | --- | --- | --- | --- | --- |
|  | E13 | E28 | E31 | E34 | E21 | E57 |
| **CP673451** | 6,402 | 10,200 | 20,770 | 3,072 | 24,000 | 4,935 |
| **SN38** | 50.3 | 416 | 55.6 | 30 | 23.6 | 24 |
| **XL888** | 37.3 | 40.9 | 24.3 | 40.1 | 34 | 29.6 |

We further investigated the potential mechanism-of-action of the validated hits via STRING-Cytoscape analysis^37^ and similarity ensemble approach (SEA^38^), which showed enrichment for cell cycle regulation, gene expression, and chemokine signalling targets (Figs. 4a-c). From a potency and medicinal chemistry perspective, the Hsp90 inhibitor XL888, is the most promising candidate because it has reached clinical trials for the treatment of various solid malignancies in combination with targeted therapies^39,40^. The chemical structure of XL888 is remarkably different to that of other Hsp90 inhibitors, further increasing its value as a novel candidate drug for GBM treatment (Supplementary Fig. 2).

**Figure 4.**
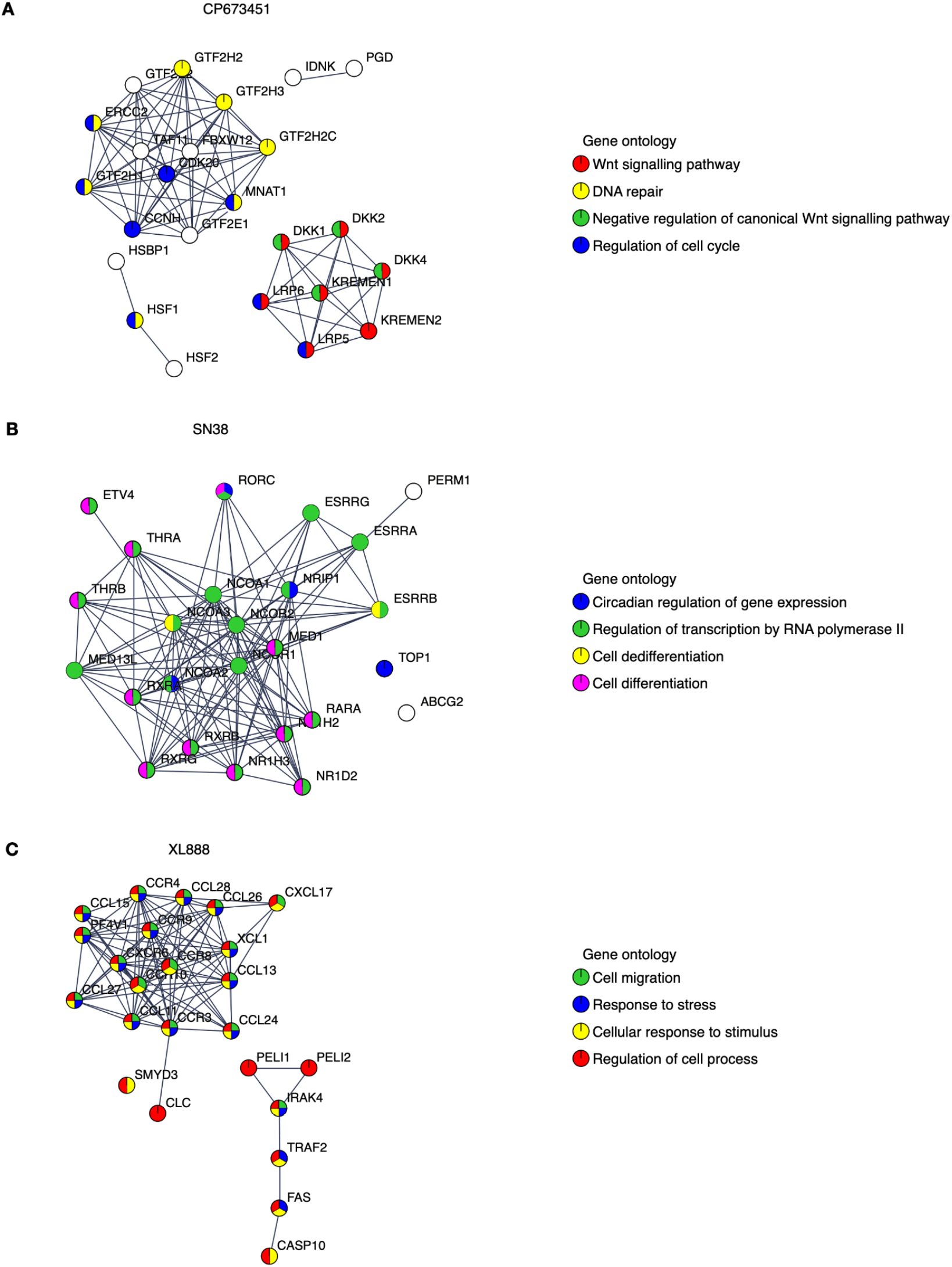
STRING pathway network analysis. (**A-C**) STRING analysis of putative intracellular pathway signalling networks influenced by each of the three compounds identified by the computational screen and characterised experimentally: **(A)** CP673451, **(B)** SN38, and **(C)** XL888. Gene ontology terms are highlighted for each network.

## Discussion

Given the high attrition rates encountered in conventional drug discovery campaigns, particularly in complex disease areas of unmet need with poorly understood targets, there is growing interest in complementary avenues that can tackle this challenge at accelerated speed. Unbiased phenotypic screening in disease relevant models that are designed to recapitulate key segments of disease pathophysiology have been proposed as a strategy to reduce late-stage attrition in drug development^41^. Drug screens performed in glioblastoma (GBM) models have so far been restricted to relatively small sets of target-annotated and approved drug libraries, limiting the potential for novel therapeutic discovery^1^. Expanding screening to larger diverse chemical libraries in complex patient-derived models quickly becomes economically infeasible to most research laboratories. Artificial Intelligence and machine learning models provide a promising route to search large areas of chemical space with relatively small amounts of data^19^.

Here, we identified and validated three compounds against the GBM phenotype, using a carefully designed prediction pipeline trained on in-house data from patient-derived GBM stem cell lines. This diverse screen captures the intrinsic heterogeneity of the GBM phenotype that led to a robust dataset for model training and a substantially higher hit identification rate than standard experimental high-throughput screening. SN38 is an active biological metabolite of the pro-drug irinotecan with primary mechanism-of-action annotated as a topoisomerase inhibitor. Topoisomerase inhibitors exhibit broad activity upon cancer cell survival^42^ and previous clinical studies have indicated that irinotecan when used in combination with other compounds might improve outcome in patients with recurrent malignant glioma^43^. However, randomized controlled clinical trials are required to fully understand the utility of irinotecan in treating GBM. Our pathway network analysis indicates SN38 may also be influencing the activity of several nuclear receptor coactivators and subfamily members such as Nuclear Receptor Subfamily 1 Group D Member 1 (NR1D1), which have previously been implicated as therapeutic targets in GBM^44^. CP673451 is a well characterized PDGFR inhibitor that has previously been demonstrated to promote GBM stem cell differentiation and inhibit GBM tumour growth *in vitro* and *in vivo*^*45*^. Our pathway network analysis indicates that CP673451 may also be potentially influencing GBM stem cell survival via modulation of heat shock factor 1 (HSF1), Kremen protein 1 (Kremen1) and General Transcription Factor IIH Subunit (GTF2H) subunit pathways. XL888 is a potent inhibitor of Hsp90 and thus exhibits a potential shared mechanism with CP673451 inhibition of HSF1 which forms a complex with Hsp90^46^. Hsp90 has previously been implicated in promoting GBM stem cell properties^47^ and Hsp90 inhibitors such as 17-AAG demonstrate inhibition of GBM growth *in vitro* and *in vivo*^*48*^. Our pathway analysis indicates XL888 may also be modulating the GBM survival phenotype via perturbation of the chemokine^49^ and tumor necrosis factor receptor-associated factor 2 (TRAF2) signalling pathways previously implicated in GBM growth and radioresistance^50^.

To the best of our knowledge, XL888 has not been previously identified as a candidate against GBM. XL888 possesses three hydrogen-bond donors and exhibits a relatively high molecular weight (503.6 Da) and cLogP (3.56), which are suboptimal features to cross the blood brain barrier. However, its modular synthesis^51^ together with its high potency against GBM cells makes XL888 a promising candidate for chemical fine-tuning into a brain-penetrant anti-GBM agent. In addition, XL888 has been shown to have protective effects on brain endothelial cells^52^, which suggests a promising safety profile. Our results therefore suggest XL888 as a candidate for further medicinal chemistry campaigns. We propose some structural changes to improve its physicochemical properties while maintaining its bioactivity (Supplementary Fig. 2, Supplementary Table 1).

## Methods

### Cell culture

Patient-derived GBM stem cell lines GCGR-E13, GCGR-E21, GCGR-E28, GCGR-E31, GCGR-E34, and GCGR-E57 were obtained from the Cancer Research UK Glioma Cellular Genetics Resource, Edinburgh. For details of the cell culture protocol for these GBM stem cell lines, please refer to Materials and Methods section from Elliott et al^24^.

### Compound screening

Cells were seeded into 384 well plates (Greiner Bio-One, uClear, 781091) and incubated for 20 hours prior to compound treatments for 72 hours. Five libraries were used for screening, giving in total 3,856 compounds. Their concentrations and number of molecules per library are shown in Fig. 1C. Out of the five channels of the original cell painting assay, only the stain for the nuclei was used in this manuscript (stain Hoechst 33342). For further details regarding this screen, please refer to Elliott et al^24^.

### Image acquisition and analysis

Plates were imaged using an ImageXpress-Confocal Ht.ai high content microscope (Molecular Devices). The images were custom-analysed with the software CellProfiler v3.1.5 (cellprofiler.org). For full details on the image acquisition and analysis, please refer to Elliott et al^24^.

### Network analysis

For the three experimentally-validated hits, we obtained a predicted target list with a similarity ensemble approach (SEA^38^). This software queries the SMILES string of an added compound to query potential compound-protein interactions across its database, taking into consideration species, Tanimoto coefficient (MaxTC) and p-value. The predicted targets were then fed into the STRING platform to assess the representation of the network components in biological pathways (KEGG database^53^) and biological processes (Gene Ontology^54^) that may be influenced by each compound.

### Data assembly for model training

To make optimal use of the compound screening data across different concentrations and heterogeneous GBM stem cell lines, we labelled compounds as positive or negative with a concentration- and cell line-dependent approach. For compounds that impacted nuclei count at only one concentration, we chose said concentration. For those that impacted nuclei count at more than one concentration, we chose the concentration in which the effect was closest to 50% nuclei count. Compounds that did not affect nuclei counts at any concentration were labelled as negative. After choosing a single concentration per compound, they were labelled as positive if they caused a decrease in nuclei count of 35% with respect to the DMSO controls in at least two, but not all, cell lines. The rest of compounds were labelled as negative. The resulting training data contained 3,458 positives and 103 negatives.

### Model training and computational screen

For model training and virtual screening, compounds were featurized with 200 physicochemical descriptors computed with the RDKit package^55^ on the SMILES strings after salt removal. The majority of SMILES were taken from the library of origin of every compound (C3L, KCGS, LOPAC, Prestwick, and Targetmol anticancer-330 for training; BioAscent and TargetMol anticancer-3,000 for screening). For chiral molecules, we employed isomeric SMILES.

Machine learning models were trained with the python libraries scikit-learn 1.3.0 and XGBoost 1.7.3. We first removed 11 features from the 200 physicochemical descriptors because they had a constant value for all compounds in the training set. Models were then trained on a reduced set of 172 *z*-score normalised features identified as relevant for classification using scikit-learn feature importance with a forest of trees function and the average reduction of Gini index as an impurity measure^56^. We retrained and tested XGBoost models on resampled 70:30 data splits, until we obtained 100 different models with test accuracy above the baseline given by the class imbalance (1 - 103/3,561= 0.971). Model hyperparameters were manually calibrated to objective = ‘binary:logistic’, learning_rate = 0.5, max_depth = 2, n_estimators = 100, and colsample_bytree = 0.5. Models were scored with precision (TP/(TP+FP)), recall (TP/(TP+FN)), F1-score (geometric mean of precision and recall), and the areas under the Receiver Operating Characteristic and Precision-Recall curves.

### Hit validation

We queried the 100 XGboost models with all compounds in the screening library to produce predicted probability of compound efficacy. Predicted hits from the computational screen were selected using a cut-off of 20% average probability across all 100 models. This led to 9/12,888 compounds in the screening library. For hit validation, we employed the same six cell lines that were used for generating the training data. Dose-responses were fitted using GraphPad Prism 9 in Fig. 3C-E for the three compounds which displayed positive action against GBM cell lines. Mean ± s.d. are shown from *n* = 3 experiments. All experimental data is available in the data source file.

## Acknowledgements

This work was supported by a Rosetrees Interdisciplinary Award (ID2022/100030) to N.O.C and D.A.O and a joint Cancer Research UK (C42454/A28596) and The Brain Tumour Charity award (GN-000676) to N.O.C. We thank Gillian Morrison from the Cancer Research UK Glioma Cellular Genetics Resource for provision of patient-derived GBM stem cell lines.

## Supplementary Material

**Supplementary Figure 1.**
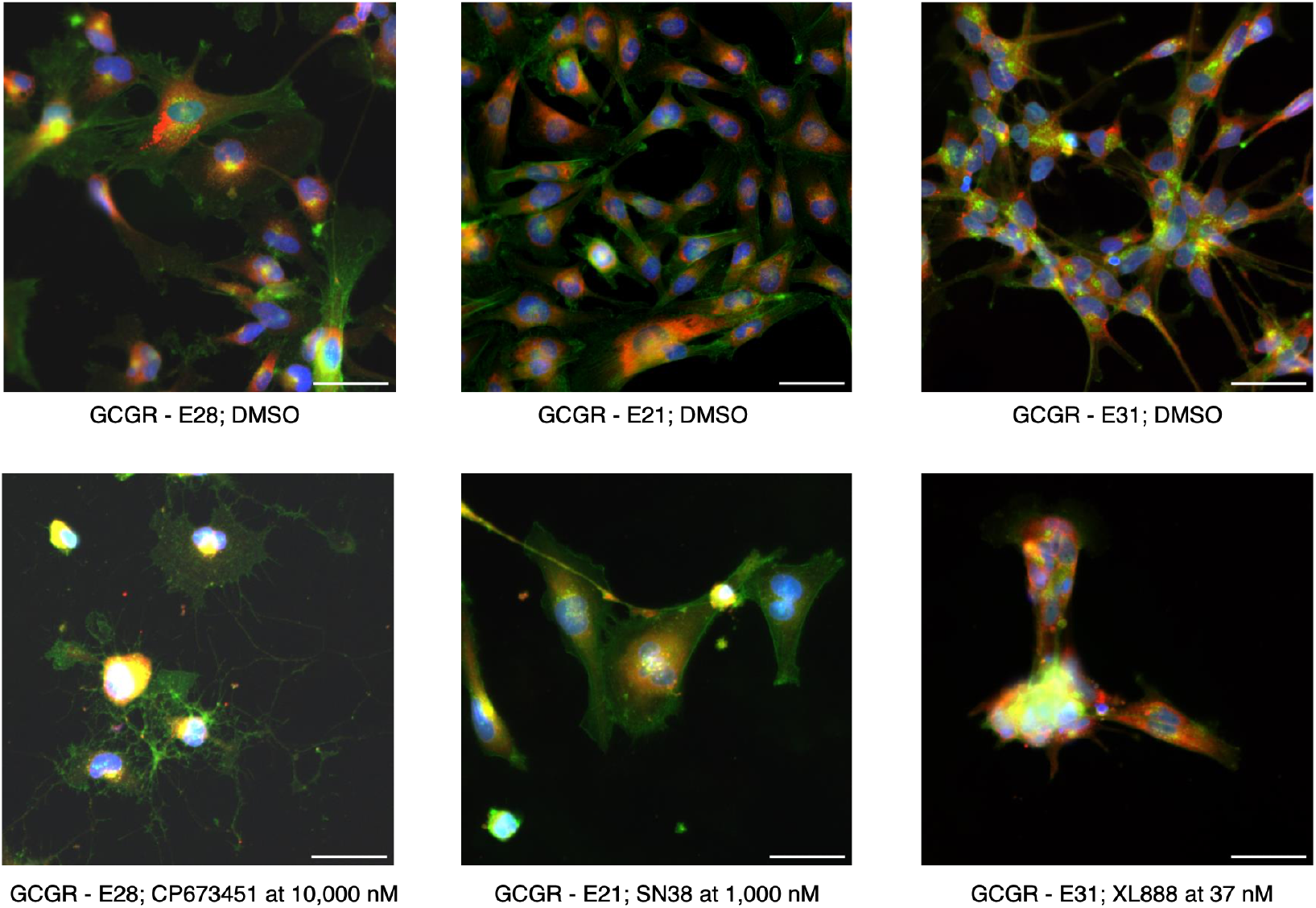
Representative images of cell line responses to compound administration vs DMSO controls. Top row: DMSO controls. Bottom row: CP673451 at 10,000 nM for cell line GCGR-E28 of classical GBM subtype, SN38 at 1,000 nM for cel line GCGR-E21 of mesenchymal GBM subtype, and XL888 at 37 nM for cell line GCGR-E31 of proneural GBM subtype.

**Supplementary Figure 2.**
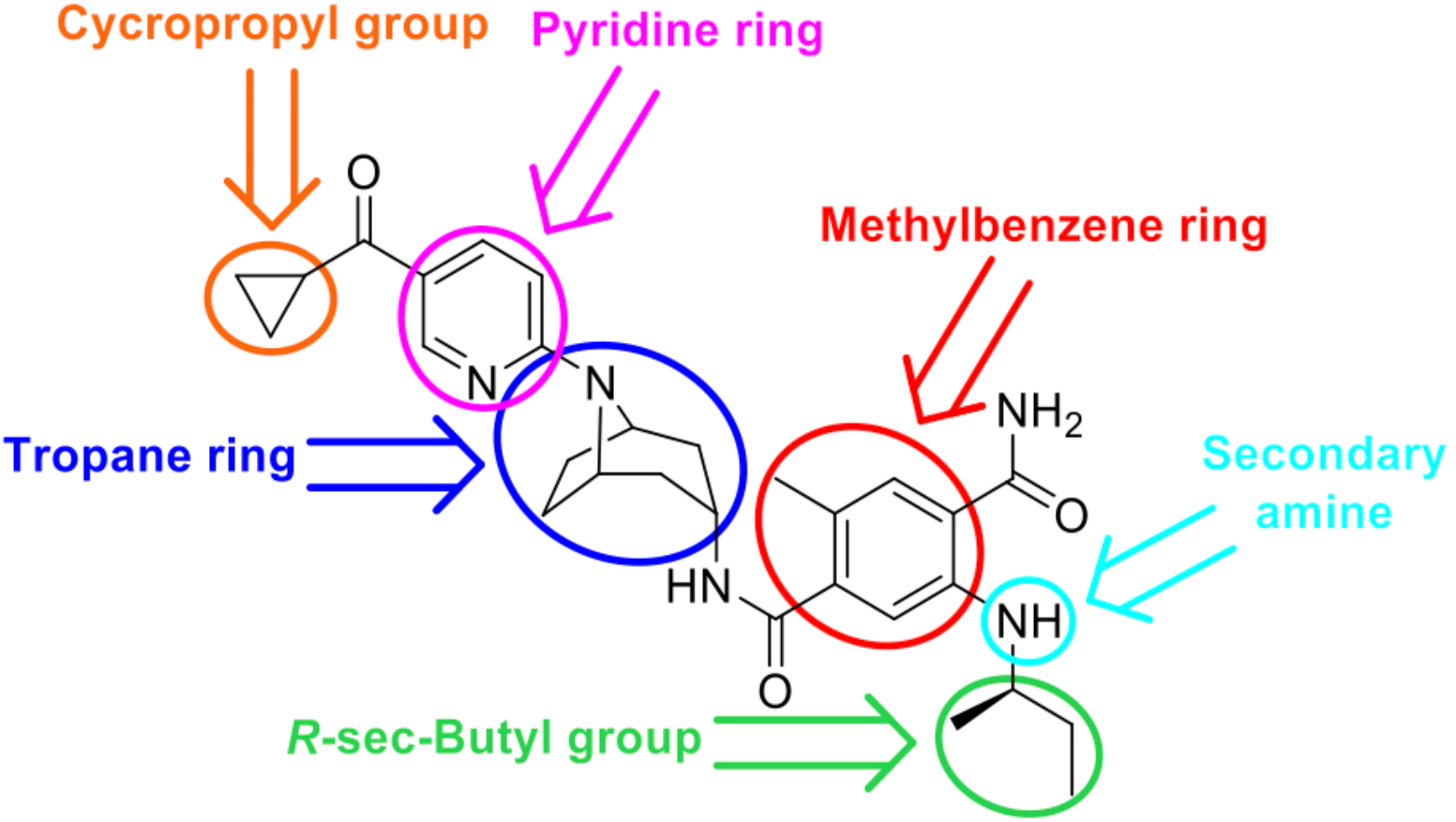
Structure of XL888. Proposed areas to modify are highlighted with coloured circles. Reduction of the size and lipophilicity of the molecule by modifying the central tropane ring, as well as the cyclopropyl and *R*-sec-butyl moieties, which are significant contributors to the overall lipophilicity of XL888. Exploration of the pyridine and methylphenyl rings by incorporating polar groups or by replacing them with bioisosteric heterocycles could also enhance BBB penetrability. Furthermore, the secondary amino group is another group that could be changed to improve the physicochemical properties of XL888. While this amino has been reported to enhance cellular activity, it does not interact with any residue of Hsp90^51^. Instead, it forms an intramolecular hydrogen bond with the carbonyl group of the benzamide ring. This interaction presents an opportunity for further medicinal chemistry optimization, potentially leading to derivatives with significantly improved BBB permeability.

**Supplementary Table 1.** Physicochemical properties and PK predictions of compound XL888.

|  |  |
| --- | --- |
| Molecular weight | 503.65 g·mol<sup>-1</sup> |
| Num. rotatable bonds | 9 |
| Num. H-bond acceptors | 6 |
| Num. H-bond donors | 3 |
| TPSA<sup>a</sup> | 117.42 Å<sup>2</sup> |
| cLogP<sup>a</sup> | 3.56 |
| Predicted solubility (pH = 7.4)<sup>b</sup> | >10 µg/mL |
| Papp (A-to-B, MDCK)<sup>c</sup> | 320 × nm/s |
| BBB penetrant<sup>a</sup> | No |
| Intestinal absorption (rat, % absorbed)<sup>c</sup> | 30% |
| CL<sub>int</sub> (rat)<sup>c</sup> | 920 mL/h/Kg |
| t<sub>1/2</sub> (rat, IV)<sup>c</sup> | 1.4 h |

<sup>a</sup> SwissADME. <sup>b</sup> DruMAP. <sup>c</sup> Bioorg. Med. Chem. Lett. 22 (2012) 5396–5404

## Notes

### Competing Interest Statement

The authors have declared no competing interest.

### Summary of Updates

The revision contains several improvements to figures and corrected typos.

